# Limited SARS-CoV-2 diversity within hosts and following passage in cell culture

**DOI:** 10.1101/2020.04.20.051011

**Authors:** Gage K. Moreno, Katarina M. Braun, Peter J. Halfmann, Trent M. Prall, Kasen K. Riemersma, Amelia K. Haj, Joseph Lalli, Kelsey R. Florek, Yoshihiro Kawaoka, Thomas C. Friedrich, David H. O’Connor

## Abstract

Since the first reports of pneumonia associated with a novel coronavirus (COVID-19) emerged in Wuhan, Hubei province, China, there have been considerable efforts to sequence the causative virus, SARS-CoV-2 (also referred to as hCoV-19) and to make viral genomic information available quickly on shared repositories. As of 30 March 2020, 7,680 consensus sequences have been shared on GISAID, the principal repository for SARS-CoV-2 genetic information. These sequences are primarily consensus sequences from clinical and passaged samples, but few reports have looked at diversity of virus populations within individual hosts or cultures. Understanding such diversity is essential to understanding viral evolutionary dynamics. Here, we characterize within-host viral diversity from a primary isolate and passaged samples, all originally deriving from an individual returning from Wuhan, China, who was diagnosed with COVID-19 and subsequently sampled in Wisconsin, United States. We use a metagenomic approach with Oxford Nanopore Technologies (ONT) GridION in combination with Illumina MiSeq to capture minor within-host frequency variants ≥1%. In a clinical swab obtained from the day of hospital presentation, we identify 15 single nucleotide variants (SNVs) ≥1% frequency, primarily located in the largest gene – ORF1a. While viral diversity is low overall, the dominant genetic signatures are likely secondary to population size changes, with some evidence for mild purifying selection throughout the genome. We see little to no evidence for positive selection or ongoing adaptation of SARS-CoV-2 within cell culture or in the primary isolate evaluated in this study.

**Author Summary:** Within-host variants are critical for addressing molecular evolution questions, identifying selective pressures imposed by vaccine-induced immunity and antiviral therapeutics, and characterizing interhost dynamics, including the stringency and character of transmission bottlenecks. Here, we sequenced SARS-CoV-2 viruses isolated from a human host and from cell culture on three distinct Vero cell lines using Illumina and ONT technologies. We show that SARS-CoV-2 consensus sequences can remain stable through at least two serial passages on Vero 76 cells, suggesting SARS-CoV-2 can be propagated in cell culture in preparation for *in-vitro* and *in-vivo* studies without dramatic alterations of its genotype. However, we emphasize the need to deep-sequence viral stocks prior to use in experiments to characterize sub-consensus diversity that may alter outcomes.

## Introduction

The emergence of severe acute respiratory syndrome coronavirus 2 (SARS-CoV-2) and coronavirus disease (COVID-19) in Wuhan, China at the end of 2019 has garnered worldwide public health attention [1–5]. At the time of writing, the United States has the highest number of confirmed cases among countries where this virus is circulating – 639,733 cases and 30,990 deaths.

The rapid spread and molecular epidemiology of SARS-CoV-2 has been tracked by sequencing viruses from infected individuals. Within weeks of the virus being identified, the complete genome was sequenced, and as of April 16th 2020, 9,330 SARS-CoV-2 genomes have been shared and used to track local transmission chains and global phylodynamics [6]. While consensus-level data has been rapidly disseminated, few researchers have analyzed viral diversity within samples below the consensus level.

SARS-CoV-2 is a betacoronavirus with 79-82% nucleotide identity shared with SARS-CoV, the virus responsible for the 2002 – 2003 SARS epidemic [7, 8]. During the 2003 SARS outbreak the virus was characterized as having gone through distinct evolutionary phases in human hosts. Initially, an excess of nonsynonymous mutations in the spike (S) gene suggested that it might be under positive selection, but this progressed into purifying selection later in the epidemic [9]. The ORF1a gene appeared to go through similar evolutionary phases as the S gene. In contrast to ORF1a and S, the ORF1b gene appeared to have undergone strong purifying selection throughout the 2003 SARS epidemic [9].

Though limited, *in-vivo* studies of SARS-CoV-2 show low-frequency variants are detectable within individual hosts and are likely due to random fluctuations in allele frequencies. One study highlights an excess of nonsynonymous variants compared to synonymous variants among these low-frequency variants, consistent with the possibility of ongoing diversifying selection in SARS-CoV-2 viruses [10, 11]. Another recent study by Liu and colleagues highlights a deletion in the Spike gene at nucleotide (nt) positions 23,585–23,599, encoding QTQTN, that flanks the polybasic cleavage site in S1/S2. The authors observe this deletion arising in SARS-CoV-2 viruses following two passages in Vero E6 cells. This deletion is found in over 50% of samples from Liu and colleagues, ranging in frequency from 8 to 33%, and is hypothesized to be adaptive for SARS-CoV-2 *in vitro,* but may be less robust *in vivo* as it was only identified in 3 of 68 Chinese-origin clinical samples at sub-consensus levels [12].

To better understand evolutionary pressures affecting SARS-CoV-2 within a single infection, we used sequence-independent, single-primer amplification (SISPA) to generate metagenomic libraries sequenced in parallel on Oxford Nanopore Technology (ONT) and Illumina sequencing platforms (**Fig 1**) [13, 14]. We obtained a nasopharyngeal (NP) swab from an individual with confirmed SARS-CoV-2 infection from the day of diagnosis, who originally presented with symptoms in Madison, WI (hereafter referred to as the Madison patient). This case was diagnosed in late January 2020 and was one of the first lab-confirmed cases in the United States. We additionally characterized viral diversity following passage in cell culture in three distinct cell types – Vero 76, Vero E6, and Vero STAT-1 knockout (KO). Passage in cell culture is expected to alter allele frequencies and may even select for adaptive mutations that make passaged viruses less representative of their genotypes and phenotypes *in vivo*. Global viral evolution ultimately derives from selective pressures and population dynamics playing out within and between individual hosts. In this study, we identify SNVs within a clinical specimen and track what happened to them through multiple rounds of passage in culture and begin to assemble a nuanced understanding of the evolution and ongoing adaptive potential of this zoonotic virus.

**Figure 1.**
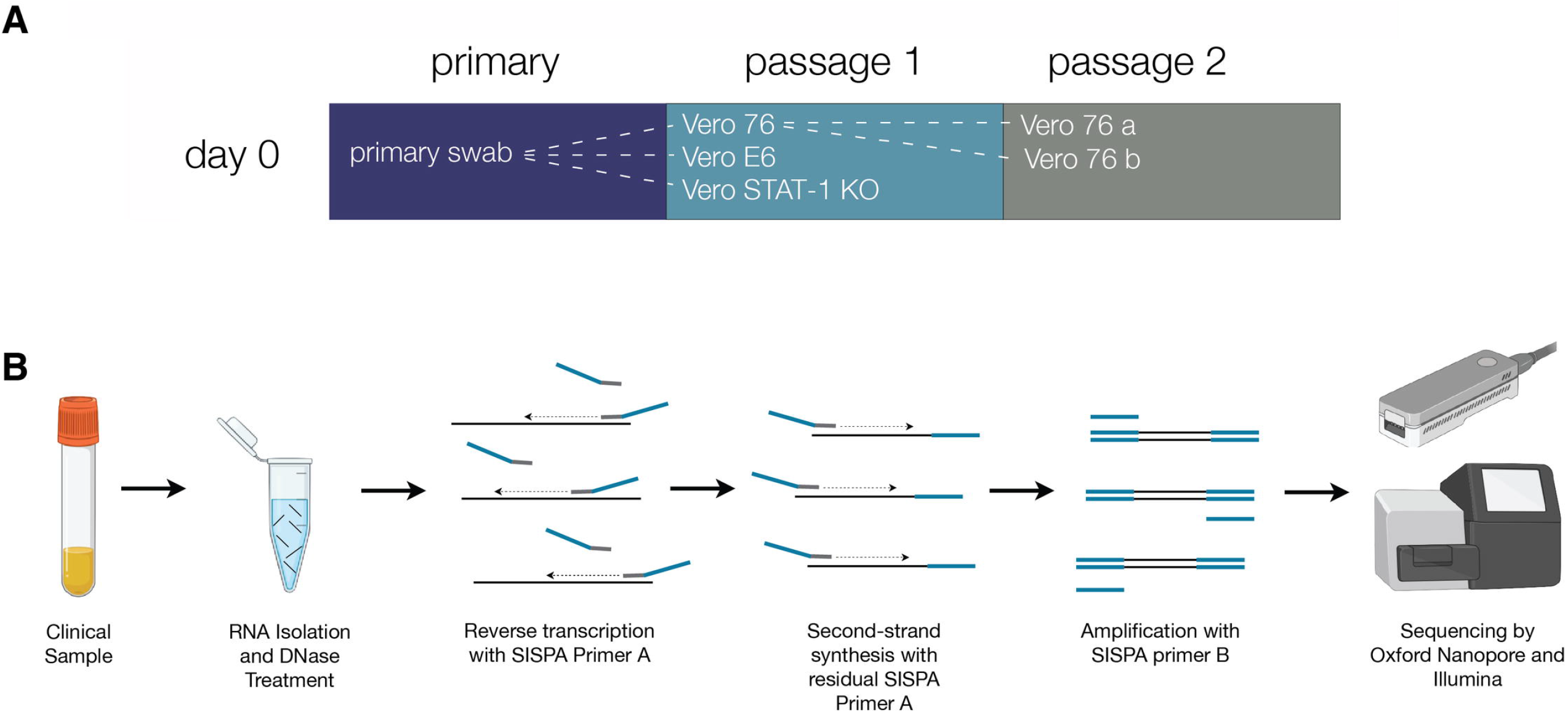
Sequence-Independent, Single-Primer Amplification sequencing workflow. A) Table showing nomenclature, and color scheme for all samples used in this study. B) Schematic showing the sequence-independent, single-primer amplification sequencing workflow.

## Results

### No consensus-level changes following two passages on Vero 76 cells

We obtained an NP swab from the day of diagnosis and passaged the virus on three distinct cell lines – Vero 76, Vero E6, and Vero STAT-1 KO (**Fig 1**). To understand the effects of serial passaging on SARS-CoV-2, we used the SISPA approach to generate full genome sequencing libraries from the original NP swab and passaged virus (**S1 Fig**). Sequences were analyzed in parallel using custom in-house scripts to deplete host reads, map to the SARS-CoV-2 Madison reference (Genbank: MT039887.1; originally sequenced by the US Centers for Disease Control and Prevention), and call minor variants ≥10% and ≥1% for ONT and Illumina datasets, respectively. We detect no consensus-changing SNVs through two passages on Vero 76 cells and through one passage on Vero E6 and Vero STAT-1 KO cells (passage 2 samples were not available in these cell lines) (**Fig 2a**).

**Figure 2.**
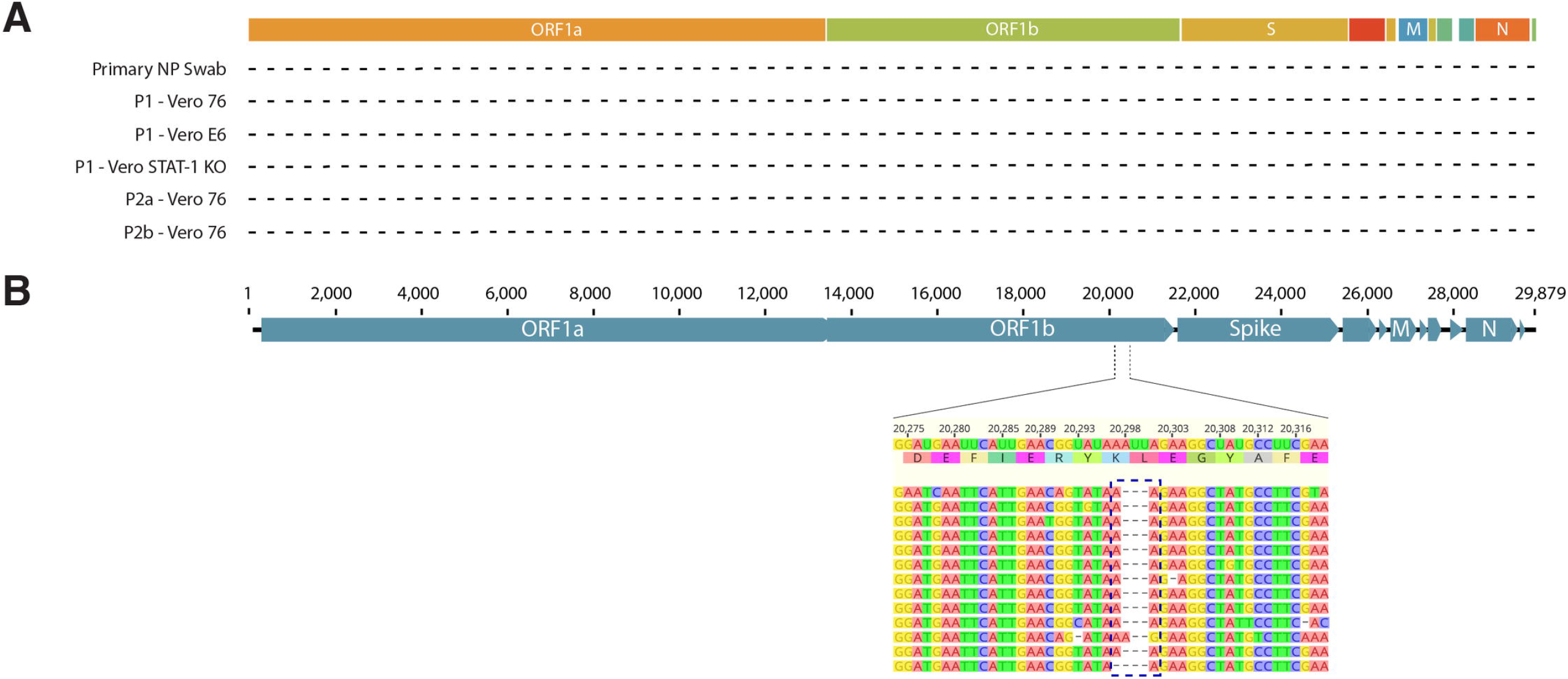
Consensus sequence overview for SARS-CoV-2 samples. A) Map of the SARS-CoV-2 genome illustrating no consensus-level changes compared to the reference (MT039887.1). B) Map of the Madison SARS-CoV-2 showing an in-frame deletion at nucleotide position 20,298 – 20,300 relative to the Wuhan reference (MN908947.3).

Interestingly, in comparison to the sequence derived from the first case of SARS-CoV-2 (MN908947.3), the Madison patient’s virus contained an in-frame deletion at nucleotide positions 20,298 – 20,300 (**Fig 2b**). This deletion has not been identified in any other samples submitted to GISAID as of April 8, 2020. This deletion occurs in a region that codes for the poly(U)-specific endoribonuclease, but its functional impact is not clear [12].

### No deletion in spike gene after passaging in cell culture

To understand how serial passaging SARS-CoV-2 affects genomic variation, we sequenced virus populations after each passage using the same SISPA metagenomics approach we used to characterize the original biological specimen. Passaged sample names and cell lines are described in the methods. An in-house pipeline (available at: https://github.com/katarinabraun/SARSCoV2_passage_MS) was applied to trim out primer sequences, bioinformatically deplete host reads, and generate alignment files, which contained all reads mapping to the SARS-CoV-2 Madison reference genome (MT039887.1). At the consensus level, SARS-CoV-2 does not accumulate genetic variation after two passages on Vero 76 cells (**Fig 2**). We also examined deletions ≥1% frequency and ≥3 nt in length. We found no evidence of deletions that fit these criteria in any of the cell culture isolates.

### Most minor variants are found in the largest genes – ORF1a and ORF1b

To characterize patterns of sub-consensus diversity, we looked at SNVs at or above 1% frequency in only the Illumina reads. We previously established that this conservative cutoff ensures that only bona fide mutations are considered [15, 16]. All minor variant analyses and figures were completed using the Illumina SNV data as these data are higher average quality and ideal for analysis involving low-frequency variants (**Fig 3**). Seventy-five percent of all minor variants we identify fall in ORF1a and ORF1b, which together take up 72.8% of the length of the 28kb coding genome. ORF1a and ORF1b encode the replicase machinery [7]. We account for differences in gene size by normalizing variants to kilobase gene length (variants / kb-gene-length – “v/kbgl”) [10]. The highest density of variants was reported in smaller genes like envelope, ORF7a, and ORF10 (**S2 Table**). We also show that through each passage, variant density in ORF1a and ORF1b increases. There were no SNVs ≥1% in the spike gene in the primary NP swab, but low-frequency SNVs (all <5%) were identified in spike following passage in cell culture (**Fig 3**). Outside of ORF1a and ORF1b, the other genes in the primary NP swab are clonal above the 1% threshold, with the exception of one low-frequency SNV in nucleoprotein (N).

**Figure 3.**
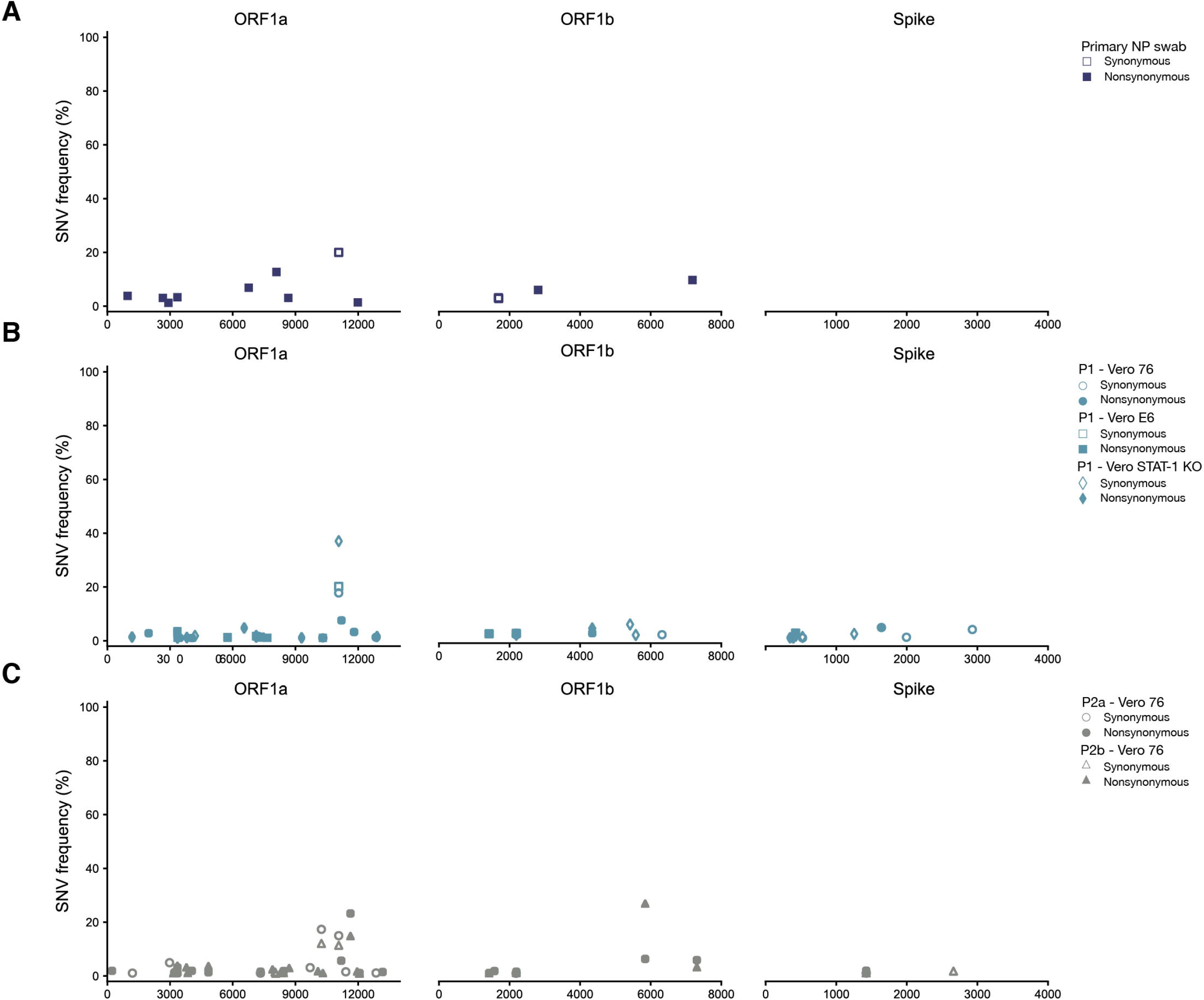
Minor variant frequencies in ORF1a, ORF1b, and Spike coding regions of the SARS-CoV-2 genome. A) Minor variants ≥1% frequency that were detected in the original primary NP swab by Illumina sequencing in ORF1a, ORF1b, and spike genes. B) Minor variants ≥1% frequency that were detected in the first passage by Illumina sequencing in ORF1a, ORF1b, and spike genes. C) Minor variants ≥1% frequency that were detected in the second passage by Illumina sequencing in ORF1a, ORF1b, and spike genes.

A few SNVs at intermediate frequencies or identified across multiple samples stood out. A synonymous SNV at nucleotide position 11,070 (ORF1a_11070_syn) was found at ≥15% frequency in the primary NP swab as well as in all passaged samples. Amino acid positions 3,570 – 3,859 in ORF1a are predicted to be involved in the formation of double-membraned vesicles [7]. Variants at nucleotide positions 127 (nonsynonymous – asparagine to aspartic acid; ORF7a_127_N43D) and 129 (synonymous; ORF7a_129_syn) were identified between 1-4% frequency in all passaged samples, but were not detected in the primary NP swab. ORF7a has no known function, so the impact of these SNVs is unclear [7]. These SNVs (ORF1a_11070_syn, ORF7a_127_N43D, and ORF7a_129_syn) have not been identified as major variants in any of the SARS-CoV-2 genomes submitted to GISAID as of 12 April, 2020. Six variants identified in at least one sample evaluated here have been identified as major variants in at least one sequence on Nextstrain as of 12 April, 2020. These SNVs include ORF1a_8025_syn (p2b Vero 76) found in England/201380056/2020, England/20146004904/2020, and Australia/VIC164/2020; ORF1a_11409_syn (p2a Vero 76) found in HongKong/HKPU2_1801/2020; ORF1b_5843_T1948I (p2a Vero 76 and p2b Vero 76) found in China/IQTC02/2020; S_1640_T547I (p1 Vero 76) found in USA/WA-S17/2020; S_2661_syn (p2b Vero 76) found in HongKong/HKPU1_2101/2020; and ORF3a_385_L129F (p1 Vero-1 STAT KO and p1 Vero 76) found in Algeria/G0638_2264/2020. Interestingly, all six of these SNVs are a cytosine to thymine transitions.

We also determined whether SNVs were shared among the primary NP swab and passaged viruses (**S2 Fig)**. Thirteen of the 15 minor variants identified in the primary NP swab are purged following passage in cell culture. Only two SNVs were found in all of the available samples – ORF1a V1118A and the synonymous SNV at nt 11,070 in ORF1a. ORF1a V1118A remains between 1-2% in all viruses. However, ORF1a 11,070-syn is found at 3% in the primary NP swab and increases in frequency to 18% in p1 Vero 76, remaining above 10% in both p2 Vero 76 samples. Only two *de novo* SNVs are found above 10% in cell culture – ORF1a_10242_syn (p2a Vero 76 and p2b Vero 76) and ORFb_5843_T1948I (p2b Vero 76).

### SNV frequency spectra reveal an excess of low-frequency SNVs

Purifying selection is known to remove new variants from the population, generating an excess of low-frequency variants, while positive and/or diversifying selection promotes the accumulation of intermediate- and high-frequency variation [17]. Especially in the setting of an acute viral infection, exponential population growth can also result in an excess of low-frequency variants. Population bottlenecks, for example sharp reductions in a viral population size typically associated with airborne viral transmission, can contribute to an excess of intermediate- and high-frequency variation. We generated site frequency spectra to expand our assessment of the evolutionary pressures impacting SARS-CoV-2 viruses within humans and in cell culture. A “neutral model” (assumes a constant population size and the absence of selection), represented in light grey in **Fig 4**, predicts around 50% of polymorphisms will be low-frequency (1-10%). In stark contrast to the neutral expectation, we observed ≥80% of SNVs falling into the low-frequency bin in the primary nasal swab sample as well as passaged samples. This dramatic excess of low-frequency variation is consistent with purifying selection acting to purge new, deleterious mutations. This signature is also consistent with population expansion as is expected in humans following airborne transmission and in cell culture after each passage.

**Figure 4.**
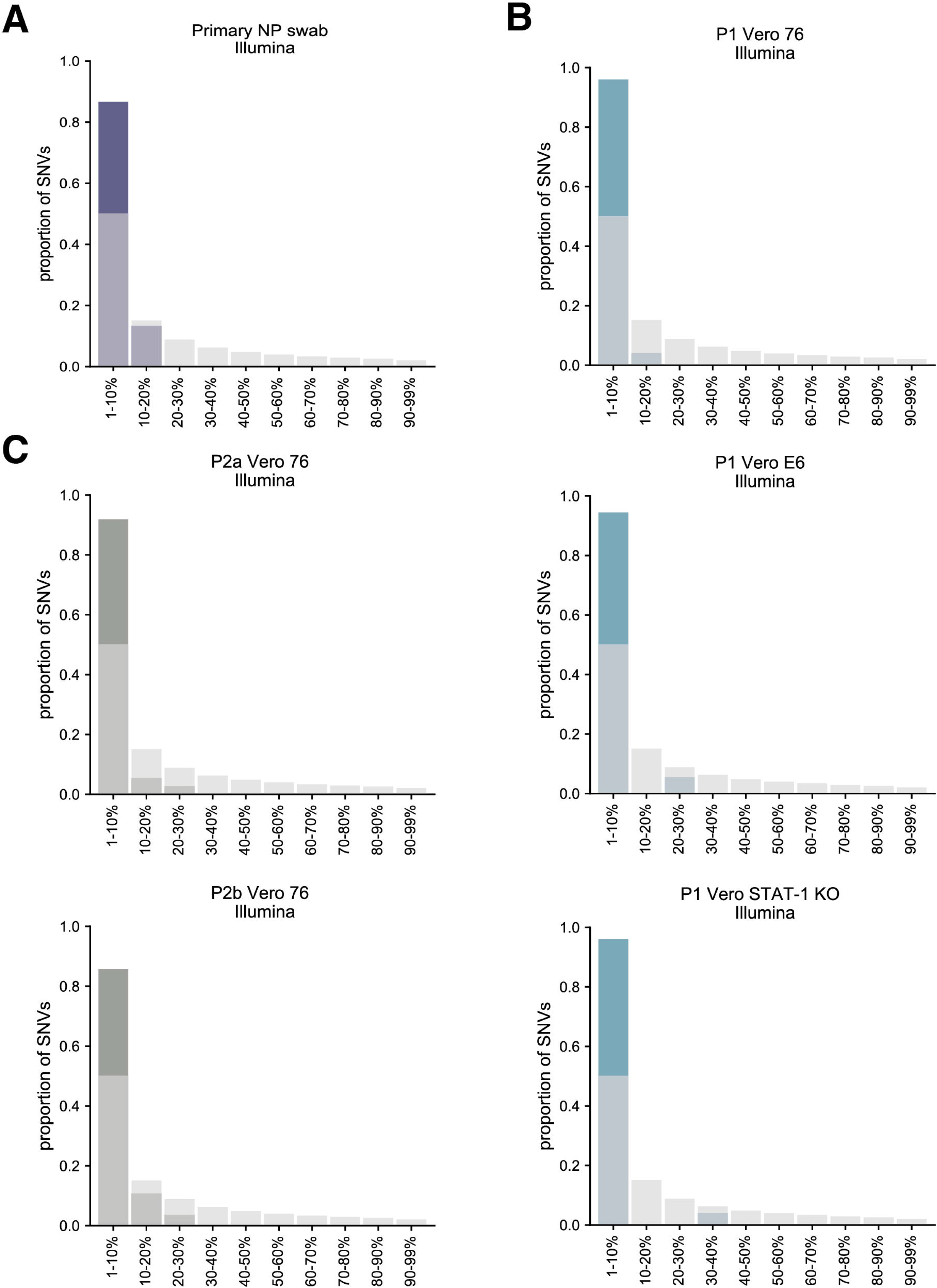
SNV frequency distributions. The frequency of Illumina detected SNVs plotted against a “neutral model”, represented in light grey. The neutral model assumes a constant population size and the absence of selection. A) SNV frequency spectrum from the primary NP swab, represented in dark blue. B) SNV frequency spectrum from three p1 samples, represented in turquoise. C) SNV frequency spectrum from two p2 samples, represented in dark grey.

### Nucleotide diversity patterns point toward mild purifying selection

In addition to assessing the fate of individual minor variants, we were also interested in evaluating population dynamics using diversity metrics. Specifically, we calculated genewise diversity using π, the average number of pairwise differences per nucleotide site among a set of sequences, for each gene in each sample. Overall, genewise nucleotide diversity is very low compared to other RNA viruses, consistent with low mutation rates in coronaviruses due to RNA proofreading machinery [18, 19]. Genewise diversity was very low in the primary NP swab and was only measurable in ORF1a (9 SNVs), ORF1b (5 SNVs) and N (1 SNV). Genewise diversity is more varied in the passaged samples (**Fig 5**). Interestingly, π is highest in ORF7a in these samples – although this signal seems to be primarily driven by the small size of this gene. To more directly assess whether SARS-CoV-2 viruses are under selective pressure in the human infection evaluated here and in cell culture, we also compared the relative abundance of nonsynonymous (πN) and synonymous (πS) polymorphisms in each gene, which is a common measure for selection that is also robust to variability in sequencing coverage depth [20]. The dominant genetic signature when looking across the entire genome is one of purifying selection (πN/πS < 1). In ORF1a, πS > πN in the primary NP swab as well as p1 and p2 samples. In ORF1b, πN/πS is close to 1 in the primary NP swab and the p1 on Vero 76 and Vero E6 cells, suggesting a more prominent role of genetic drift in this gene. Interestingly, πN/πS >> 1 in p1 Vero 76 ORF10, p1 Vero E6 envelope (E), and p1 Vero STAT-1 KO ORF3a.

**Figure 5:**
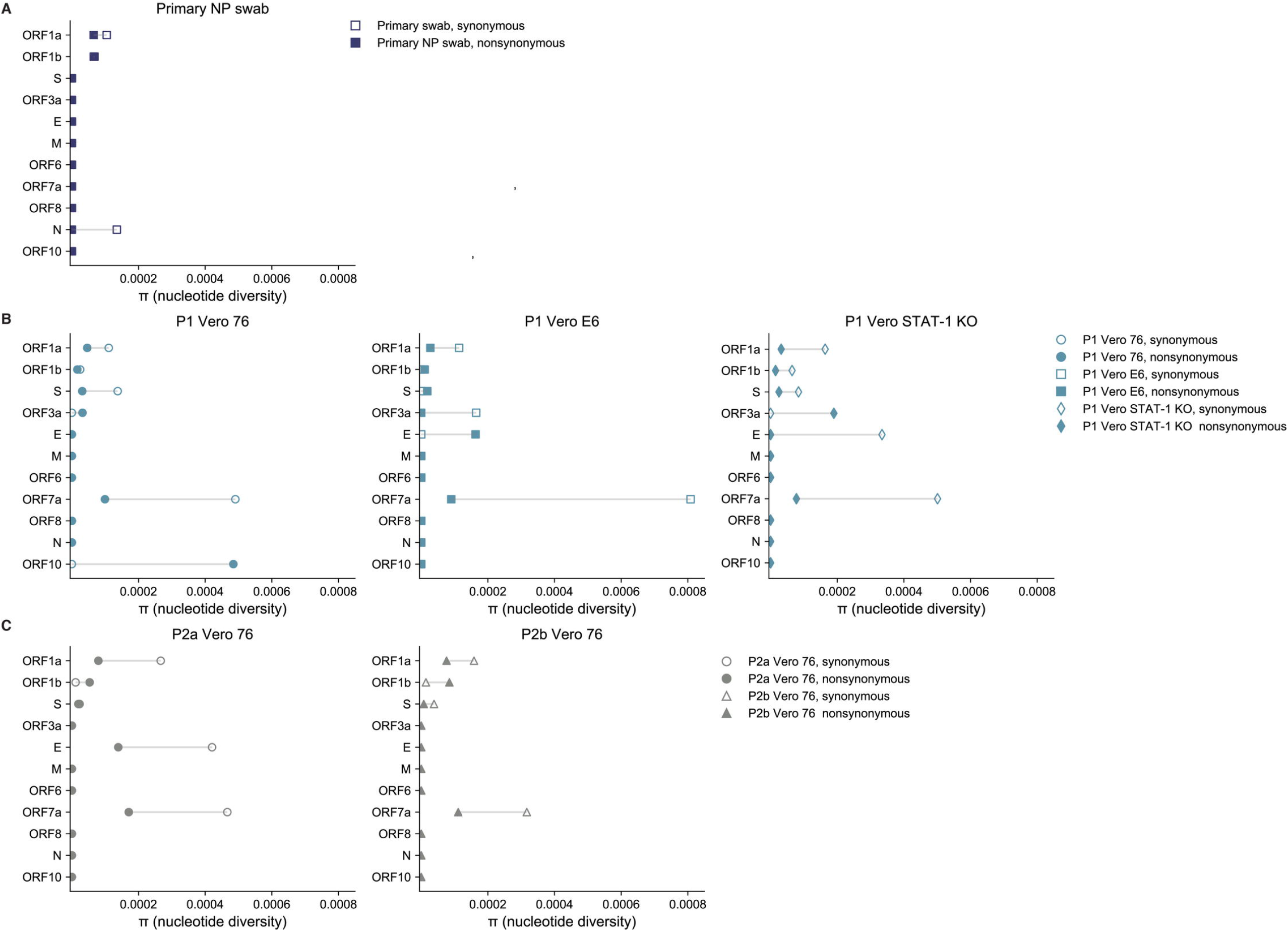
Intragene nucleotide diversity. Relative abundance of nonsynonymous (πN) and synonymous (πS) for all 11 open reading frames. Nonsynonymous diversity (πN) is denoted by closed symbols and synonymous diversity (πS) is denoted by open symbols. A) Intragene π from the primary NP swab, represented in dark blue. B) Intragene π from three p1 samples, represented in turquoise. C) Intragene π from two p2 samples, represented in dark grey. Length of horizontal line is the difference between πN and πS for each gene.

### Comparison of Illumina and ONT ability to capture minor variant frequencies

We examined the concordance between SNV calls at the same sites, irrespective of frequency, determined by Illumina and ONT workflows. To begin, we used a stringent cutoff of 10% frequency for ONT SNVs. We then called variants at percentage frequencies decreasing by 0.5% (eg. calling 8% variants, then 7.5%, etc) until the variants called by ONT no longer matched Illumina variants irrespective of frequency at these sites (**Fig 6**, **S1 Table.**). We found that for the primary NP swab we were able to call minor variants that occurred at ≥8% frequency. Below 8% frequency, SNVs called by ONT were no longer exactly concordant with SNVs called by Illumina. Discrepancies between ONT and Illumina variant calls at low frequencies are tied to ONT’s high false discovery rate, a finding previously documented by Grubaugh and colleagues in 2019 [21]. For the p1 samples, ONT was able to capture variants that occurred at ≥4.5% frequency. For the p2 samples, we called SNVs down to 8.5% and 5.5% for the p2a and p2b samples, respectively. We likely observed concordant SNV calls between Illumina and ONT at lower frequencies in the passaged samples because viral titer *in vitro* typically exceeds viral titer *in vivo* resulting in higher average coverage in the passaged samples required to support minor variant calls at lower frequencies.

**Figure 6.**
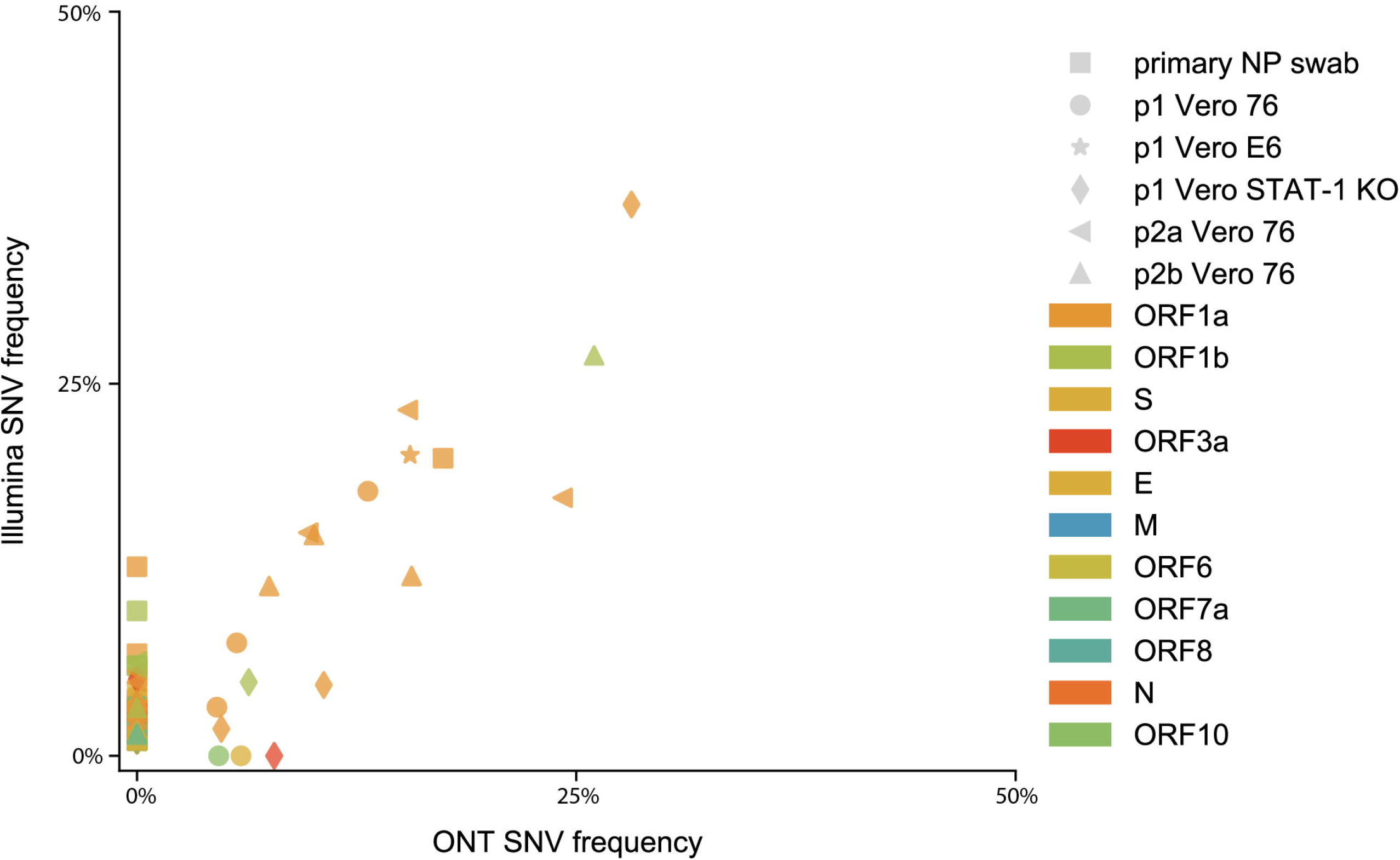
Comparison of ONT and Illumina SNV calls. Concordance between SNV calls at the same sites, irrespective of frequency, determined by Illumina and ONT workflows. Symbol denotes sample and color denotes gene. Gene colors correspond to the genome map in Figure 2.

## Discussion

Minor variants are critical for addressing molecular evolution questions, identifying selective pressures imposed by vaccine-induced immunity and antiviral therapeutics, and characterizing interhost dynamics, including the stringency and character of transmission bottlenecks. Parallel consensus-level data of clinical isolates are similarly important and allow us to predict transmission patterns on a global, regional, and community-wide scale. Here, we explore SARS-CoV-2 intrahost variation from a primary NP swab as well as from viruses passaged on three distinct Vero cell lines. We show that while diversity is low overall, the dominant viral genetic signature is one of mild purifying selection, evidenced by an excess of low-frequency variants and the observation that πN/πS < 1 in most genes across all samples evaluated.

We show that SARS-CoV-2 consensus sequences can remain stable through at least two serial passages on Vero 76 cells even in the presence of a three nucleotide deletion in the region of the genome encoding the poly(U)-specific endoribonuclease, suggesting SARS-CoV-2 can be propagated in cell culture in preparation for *in vitro* and *in vivo* studies without dramatic alterations of its genotype. A recent paper by Duggal et al. illustrate the importance of viral genotype instability in Zika virus (ZIKV) by describing variants enriched during cell culture passage (Envelope-330L/NS1-98G), despite being attenuated *in vivo* and responsible for a less pathogenic phenotype in mice compared to the wildtype genotype (Envelope-330V/NS1-98W) [22]. Viral genotype instability in cell culture can significantly affect animal model development and vaccine efficacy studies.

Though we do detect a handful of minor variants in ORF1a and ORF1b in the primary NP swab, it is notable that eight out of eleven genes are clonal above the 1% frequency level. As natural selection can only act upon genetic variation already existing within a population, very limited intrahost genetic diversity suggests the pace of SARS-CoV-2 evolution may be primarily limited by the generation of *de novo* variants. It is unclear at this time the degree to which limited within-host viral diversity is linked to coronavirus biology – e.g. RNA proofreading capabilities, homologous recombination allowing for the decoupling of deleterious “hitchhiker” mutations, and a comparatively low mutation rate. Studies have estimated the mutation rate of coronaviruses to be 2 × 10^−6^ mutations per site per round of replication, which is in line with other coronaviruses [18], but lower than influenza, 7.1 × 10^−6^ − 4.5 × 10^−5^ mutations per site per round of replication, another respiratory RNA virus [23–27].

A previous study claimed that a common deletion at nt position 23,585–23,599 (spike), encoding QTQTN, arises after two passages in Vero E6 cells [12]. We did not identify similar deletions in this region in any of our passaged samples, suggesting this deletion is not as common as previously suggested. Interestingly, the primary NP swab obtained from the Madison patient on the day of diagnosis contained an in-frame deletion at nucleotide positions 20,298 – 20,300 (ORF1ab) that was retained through two passages on Vero 76 cells. These genomic deletions highlight the importance of characterizing viral stocks by deep-sequencing so genotypic differences that may alter experimental outcomes can be thoroughly documented and shared with other researchers.

Below the consensus level, we found an excess of low-frequency variants compared to what would be expected in a neutral setting with no changes in population size and no selective pressures at play. This suggests that either purifying selection is acting to remove new, mildly deleterious mutations in hosts and in culture before they can reach intermediate or high frequencies, and/or the virus is undergoing exponential population growth as would be expected in an acute viral infection or following passage in cell culture. It is likely that viral exponential population growth is contributing to this genetic signature; however, without additional samples, it is difficult to determine the relative contribution of each of these factors. We would emphasize these findings are rooted in relatively few, low-frequency SNVs from a single time point so conclusions about the overall evolution of SARS-CoV-2 are necessarily limited. Continued deep sequencing and analyses of SARS-CoV-2 minor variant SNV populations in humans and in cell culture are critical.

## Methods

### Sample collection and cell culture passage conditions

Three different Vero cell lines were purchased from ATCC; Vero 76 (ATCC: CRL-1587), Vero C1008 (ATCC: CRL-1586), Vero STAT-1 KO (ATCC: CCL-81-VHG), and were grown in Minimum Essential Medium (MEM) supplemented with 10% fetal bovine serum (FBS) and L-glutamine at 37°C with 5% CO_2_.

For the initial infection, the original clinical nasopharyngeal (NP) swab was divided evenly between three TC25 cm^2^ flasks seeded the day before with 1 × 10^6^ cells per flask; one flask for each Vero cell line. Virus in the original clinical sample was layered onto the cells for one hour at 37°C, the flasks were washed once with MEM, and the medium was replaced with fresh MEM supplemented with 2% FBS. For each additional passage, cells were seeded in 75 cm^2^ flasks the day before infection with 4 × 10^6^ cells per flask and infected at a multiplicity of infection between 0.01-0.001. For each passage, the virus was harvested when cell death was observed to be around 80% (~4-5 days after infection).

Work with live virus was performed at biosafety level-3 containment at the Influenza Research Institute at the University of Wisconsin – Madison under a recombinant DNA protocol approved by the Institutional Biosafety Committee. Approval to obtain the de-identified clinical sample was reviewed by the Human Subjects Institutional Review Boards at the University of Wisconsin – Madison.

### Nucleic acid extraction

For each sample, approximately 140 μL of viral transport medium or cell culture supernatant was passed through a 0.22μm filter (Dot Scientific, Burton, MI, USA). Total nucleic acid was extracted using the Qiagen QIAamp Viral RNA Mini Kit (Qiagen, Hilden, Germany), substituting carrier RNA with linear polyacrylamide (Invitrogen, Carlsbad, CA, USA) and eluting in 30 μL of nuclease free H_2_O. Samples were treated with TURBO DNase (Thermo Fisher Scientific, Waltham, MA, USA) at 37°C for 30 min and concentrated to 8μL using the RNA Clean & Concentrator-5 kit (Zymo Research, Irvine, CA, USA). Full protocol for nucleic acid extraction and subsequent cDNA generation is available at https://www.protocols.io/view/sequence-independent-single-primer-amplification-o-bckxiuxn.

### Complementary DNA (cDNA) generation

Complementary DNA (cDNA) was synthesized using a modified Sequence Independent Single Primer Amplification (SISPA) approach described by Kafetzopoulou et al. [14]. RNA was reverse transcribed with SuperScript IV Reverse Transcriptase (Invitrogen, Carlsbad, CA, USA) using Primer A (5’-GTT TCC CAC TGG AGG ATA-(N_9_)-3’). Reaction conditions were as follows: 1μL of primer A was added to 4 μL of sample RNA, heated to 65°C for 5 minutes, then cooled to 4 ◻ for 5 minutes. Then 5 μL of a master mix (2 μL 5x RT buffer, 1 μL 10 mM dNTP, 1 μL nuclease free H_2_O, 0.5 μL 0.1M DTT, and 0.5 μL SSIV RT) was added and incubated at 42◻ for 10 minutes. For generation of second strand cDNA, 5 μL of Sequenase reaction mix (1 μL 5x Sequenase reaction buffer, 3.85 μL nuclease free H_2_O, 0.15 μL Sequenase enzyme) was added to the reaction mix and incubated at 37°C for 8 minutes. This was followed by the addition of a secondary Sequenase reaction mix (0.45 μl Sequenase Dilution Buffer, 0.15 μl Sequenase Enzyme), and another incubation at 37◻ for 8 minutes. Following incubation, 1μL of RNase H (New England BioLabs, Ipswich, MA, USA) was added to the reaction and incubated at 37°C for 20 min. Conditions for amplifying Primer-A labeled cDNA were as follows: 5 μL of primer-A labeled cDNA was added to 45 μL of AccuTaq master mix per sample (5 μL AccuTaq LA 10x Buffer, 2.5 μL dNTP mix, 1μL DMSO, 0.5 μL AccuTaq LA DNA Polymerase, 35 μL nuclease free water, and 1 μL Primer B (5′-GTT TCC CAC TGG AGG ATA-3′). Reaction conditions for the PCR were: 98°C for 30s, 30 cycles of 94°C for 15 s, 50°C for 20 s, and 68°C for 2 min, followed by 68°C for 10 min.

### Oxford nanopore library preparation and sequencing

Amplified cDNA was purified using a 1:1 concentration of AMPure XP beads (Beckman Coulter, Brea, CA, USA) and eluted in 48μL of water. A maximum of 1 μg of DNA was used as input into Oxford Nanopore kits SQK-LSK109. Samples were barcoded using the Oxford Nanopore Native Barcodes (EXP-NBD104 and EXP-NBD114), and pooled to a total of 140ng prior to being run on the appropriate flow cell (FLO-MIN106) using the 72hr run script.

### Nextera XT Illumina library preparation and sequencing

Amplified cDNA was purified using a 1:1 concentration of AMPure XP beads (Beckman Coulter, Brea, CA, USA) and eluted in 48μL of water. PCR products were quantified using Qubit dsDNA high-sensitivity kit (Invitrogen, USA) and were diluted to a final concentration of 0.2 ng/μl (1 ng in 5 μl volume). Each sample was then made compatible for deep sequencing using the Nextera XT DNA sample preparation kit (Illumina, USA). Specifically, each sample was enzymatically fragmented and tagged with short oligonucleotide adapters, followed by 14 cycles of PCR for template indexing. Samples were purified using two consecutive AMPure bead cleanups (0.5x and 0.7x) and were quantified once more using Qubit dsDNA high-sensitivity kit (Invitrogen, USA). The average sample fragment length and purity was determined using Agilent High Sensitivity DNA kit and the Agilent 2100 Bioanalyzer (Agilent, Santa Clara, CA). After passing quality control measures, samples were pooled equimolarly to a final concentration of 4 nM, and 5 μl of each 4 nM pool was denatured in 5 μl of 0.2 N NaOH for 5 min. Four samples (primary NP swab, p1 Vero 76, p1 Vero E6, and p1 Vero STAT-1 KO) were pooled on a single flowcell to a final concentration of 8pM with a PhiX-derived control library accounting for 1% of total DNA and was loaded onto a 500-cycle v2 flowcell. The p2 samples (p2a Vero 76 and p2b Vero 76) were pooled with seven other samples (not included in this manuscript) and were denatured to a final concentration of 14pM with a PhiX-derived control library accounting for 1% of total DNA and was loaded onto a 600-cycle v3 flowcell. Average quality metrics were recorded, reads were demultiplexed, and FASTQ files were generated on Illumina’s BaseSpace platform.

### Sequence read mapping and variant calling by ONT

Seventy-two hours after sequencing was initiated, raw sequencing reads were demultiplexed using qcat (https://github.com/nanoporetech/qcat). In order to deplete host sequences, sequencing reads are mapped against host genome and transcript references, and unmapped reads are saved. Reads were then trimmed by 30bp on each side to discard SISPA primer sequences. In this step, reads with quality scores ≤ 7 were discarded. Cleaned viral reads were then mapped to the severe acute respiratory syndrome coronavirus 2 isolate 2019-nCoV/USA-WI1/2020 consensus sequence (Genbank: MT039887.1, originally sequenced by the CDC) using minimap2. Minor variants from ONT sequences that comprise at least 10% of total sequences in any of the samples were identified using the bbmap callvariants.sh tool (https://jgi.doe.gov/data-and-tools/bbtools/). The entire ONT analysis pipeline is available at this GitHib address https://github.com/katarinabraun/SARSCoV2_passage_MS.

### Illumina sequence data analysis – quality filtering and variant calling

FASTQ files were initially processed using custom bioinformatic pipelines, available with instructions for use at the GitHub repository accompanying this manuscript https://github.com/katarinabraun/SARSCoV2_passage_MS. Briefly, read ends were trimmed to achieve an average read quality score of Q30 and a minimum read length of 100 bases using Trimmomatic (http://www.usadellab.org/cms/?page=trimmomatic) [28]. Paired-end reads were merged and then mapped to the reference sequence (Genbank MT039887.1: 2019-nCoV/USA-WI1/2020) using Bowtie2 (http://bowtie-bio.sourceforge.net/bowtie2/manual.shtml). Single nucleotide variants (SNVs) were called with Varscan2 (http://varscan.sourceforge.net/using-varscan.html) using a frequency threshold of 1%, a minimum coverage of 100 reads, and a base quality threshold of Q30 or higher [29]. SNVs were annotated to determine the impact of each variant on the amino acid sequence. SNVs were annotated in eleven open reading frames: ORF1a (open reading frame 1a), ORF1b (open reading frame 1b), S (Spike, encodes surface protein), ORF3a (open reading frame 3a), E (envelope), M (membrane), ORF6 (open reading frame 6), ORF7a (open reading frame 7a), ORF8 (open reading frame 8), N (nucleocapsid), ORF10 (open reading frame 10). VCF files were cleaned for additional analyses and figure-generation using custom Python scripts, which are all available at the GitHub repository accompanying this manuscript.

### Illumina sequence data analysis – diversity statistics

Nucleotide diversity was calculated using π summary statistics. π quantifies the average number of pairwise differences per nucleotide site among a set of sequences and was calculated per gene using SNPGenie (https://github.com/chasewnelson/SNPgenie) [30]. SNPGenie adapts the Nei and Gojobori method of estimating nucleotide diversity (π), and its synonymous (π_S_) and nonsynonymous (π_N_) partitions from next-generation sequencing data [31]. As most random nonsynonymous mutations are likely to be disadvantageous, we expect π_N_ = π_S_ indicates neutrality suggesting that allele frequencies are determined primarily by genetic drift. π_N_ < π_S_ indicates purifying selection is acting to remove new deleterious mutations, and π_N_ > π_S_ indicates diversifying selection is favoring new mutations and may indicate positive selection is acting to preserve multiple amino acid changes [32].

### Approvals

#### Biosafety

Work with live virus was performed at biosafety level-3 containment at the Influenza Research Institute at the University of Wisconsin – Madison under a recombinant DNA protocol approved by the Institutional Biosafety Committee.

#### Human subjects

Approval to obtain the de-identified clinical sample was reviewed by the Human Subjects Institutional Review Boards at the University of Wisconsin – Madison.

## Data availability

Metagenomic sequencing data after mapping to SARS-COV-2 reference genome (MT039887.1) have been deposited in the Sequence Read Archive (SRA) under bioproject PRJNA607948. Derived data, analysis pipelines, and figures have been made available for easy replication of these results at a publicly-accessible GitHub repository: https://github.com/katarinabraun/SARSCoV2_passage_MS. A description of these results is also available on LabKey at go.wisc.edu/qca2m5.

## Figure generation

Figures 3, 4, 5, 6 and supplemental figures 2 and 3 were generated using custom Python scripts and Matplotlib (https://matplotlib.org/). All code to replicate these figures can be found in the GitHub repository. Figure 1 was created with BioRender (https://biorender.com/). Supplemental figure 1 was created with JMP (https://www.jmp.com/)

## Supporting Information

### Supplemental Figure Captions

**Supplemental figure 1. Coverage depth across the SARS-CoV-2 genome.** The relative depth of coverage for each nucleotide position was plotted for (A) ONT and (B) Illumina sequencing results.

**Supplemental figure 2. Change in SNV frequency over passage.** SNVs found shared across the primary NP swab, p1 Vero 76 and p2a/p2b Vero 76 are plotted here. Symbol denotes the specific SNV. Line-type denotes route: either swab → p1 Vero 76 → p2a Vero 76 (dashed) or swab → p1 Vero 76 → p2a Vero 76 (solid). Color denotes the gene where the SNV was found. (A) Y-axis is scaled to visualize all shared SNVs, ranging from 0 – 50% frequency. (B) Y-axis is magnified to visualize SNV frequencies below 5%.

**Supplemental Figure 3. Minor frequency variants across the whole SARS-CoV-2 genome.**

### Supplemental Tables

**Supplemental Table 1.**
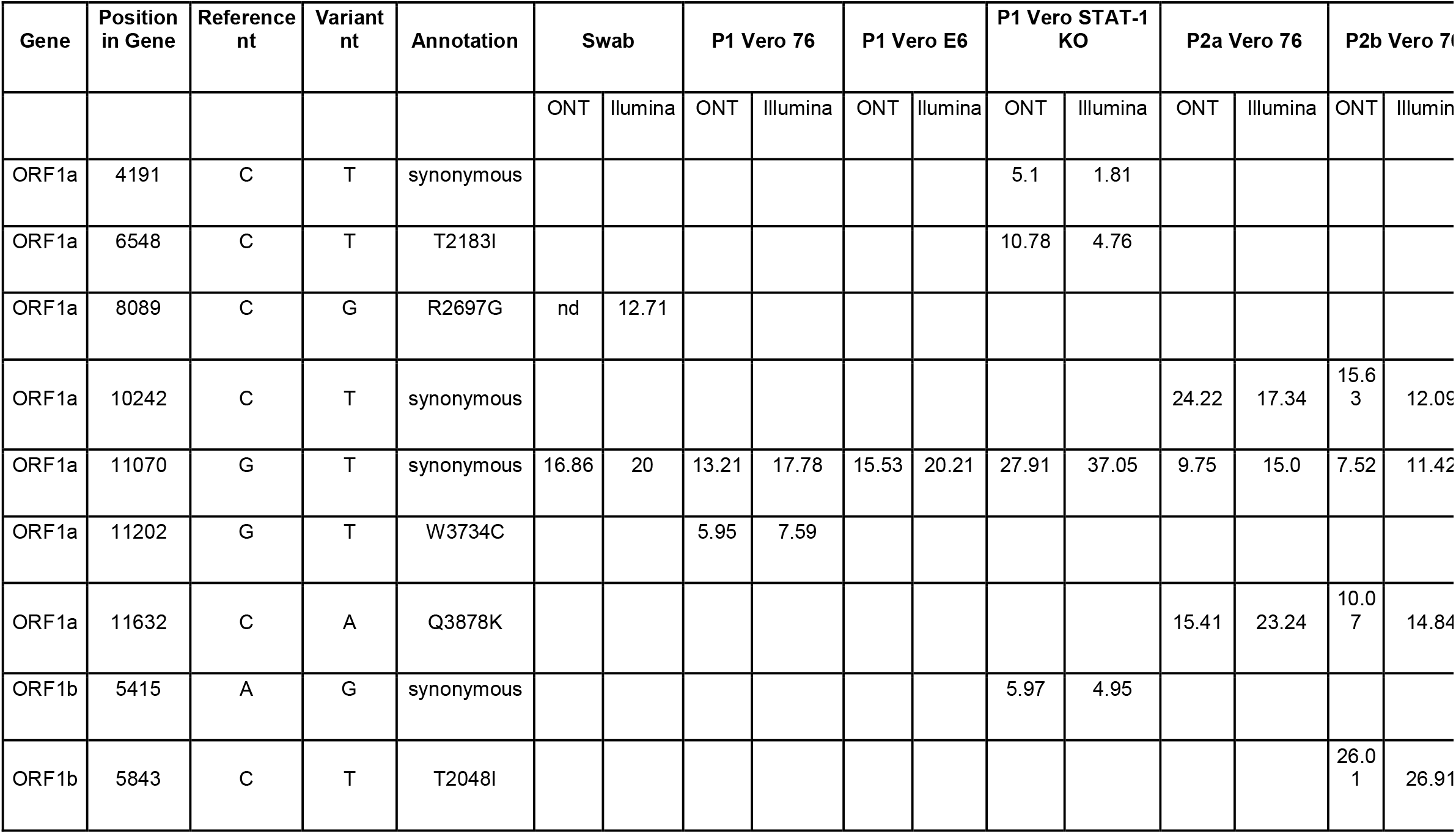
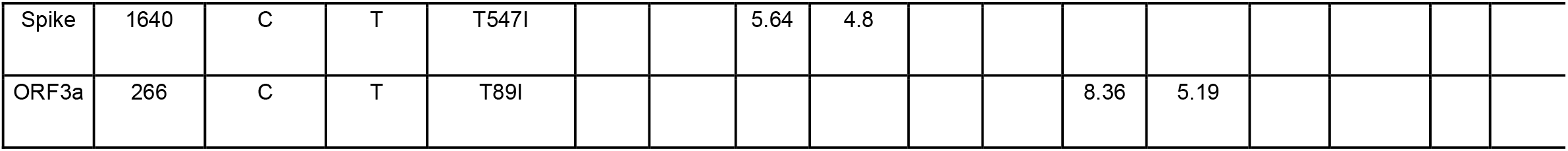
Comparison of ONT and Illumina SNVs. ‘nd’ indicates that the variant was not detected.

**Supplemental Table 2.** Variants per gene kilobase length. To normalize the number of SNVs per gene segment, we report the density of variants normalized to gene kilobase length.

| Gene | Swab | P1 Vero 76 | P1 Vero E6 | P1 Vero STAT-1 KO | P2a Vero 76 | P2b Vero 76 |
| --- | --- | --- | --- | --- | --- | --- |
| ORF1a | 0.6817 | 0.9689 | 0.8332 | 0.8332 | 1.666 | 1.439 |
| ORF1b | 0.6185 | 0.6185 | 0.2474 | 0.4948 | 0.7422 | 0.6185 |
| Spike | - | 1.0468 | 0.2617 | 1.3085 | 0.7851 | 0.5234 |
| ORF3a | - | 1.2091 | 1.2091 | 2.4183 | - | - |
| E | - | - | 4.4052 | 4.4052 | 8.8105 | - |
| M | - | - | - | - | - | - |
| ORF6 | - | - | - | - | - | - |
| ORF7a | - | 5.4794 | 5.4794 | 5.4794 | 8.2192 | 5.4794 |
| ORF8 | - | - | - | - | - | - |
| N | 0.7942 | - | - | - | 0.7942 | - |
| ORF10 | - | 8.6206 | - | - | - | - |

