## Supplementary figures and images for "Limited SARS-CoV-2 diversity within hosts and following passage in cell culture"

### Supplemental Figure 1

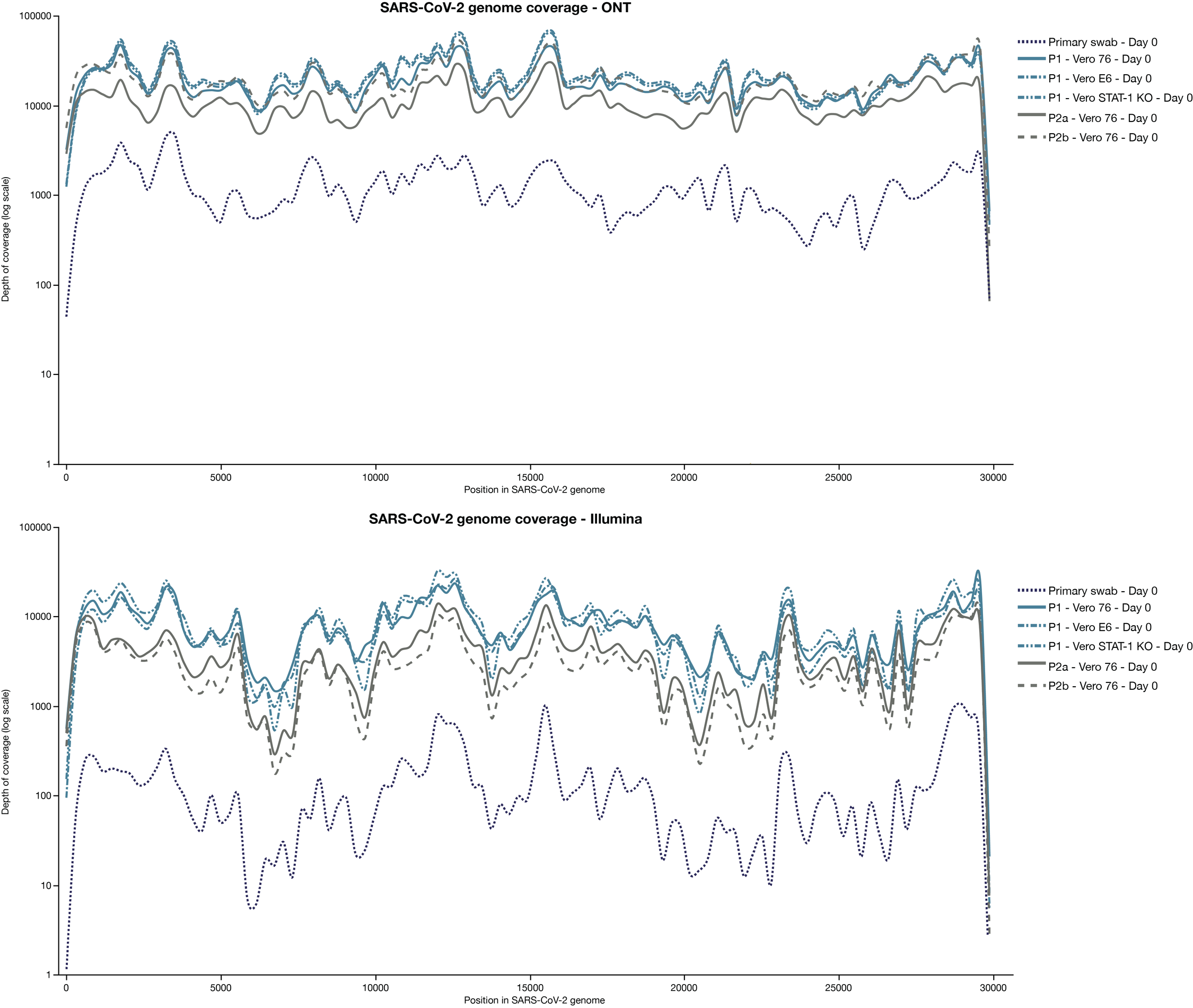

### Supplemental Figure 2

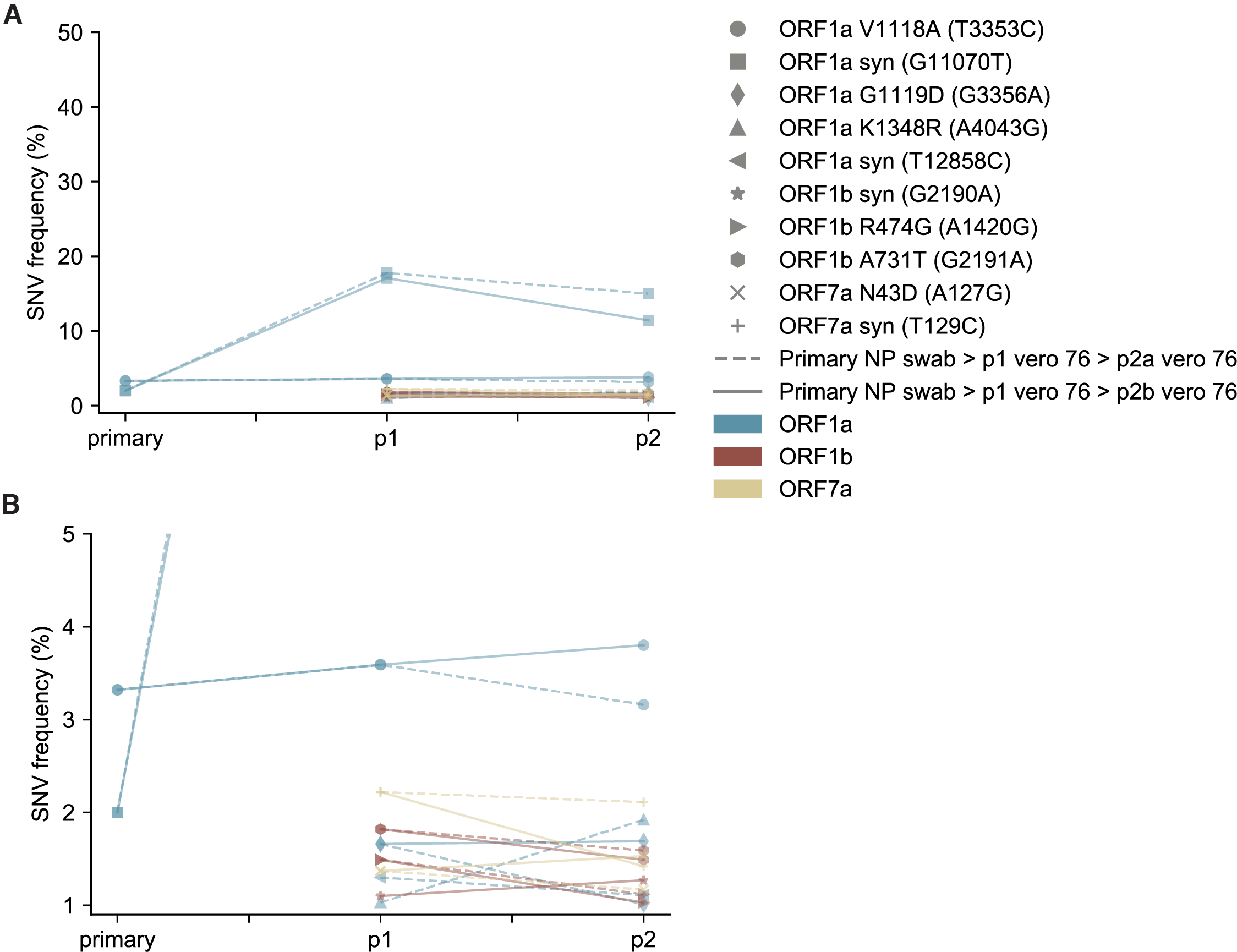

### Supplemental Figure 3

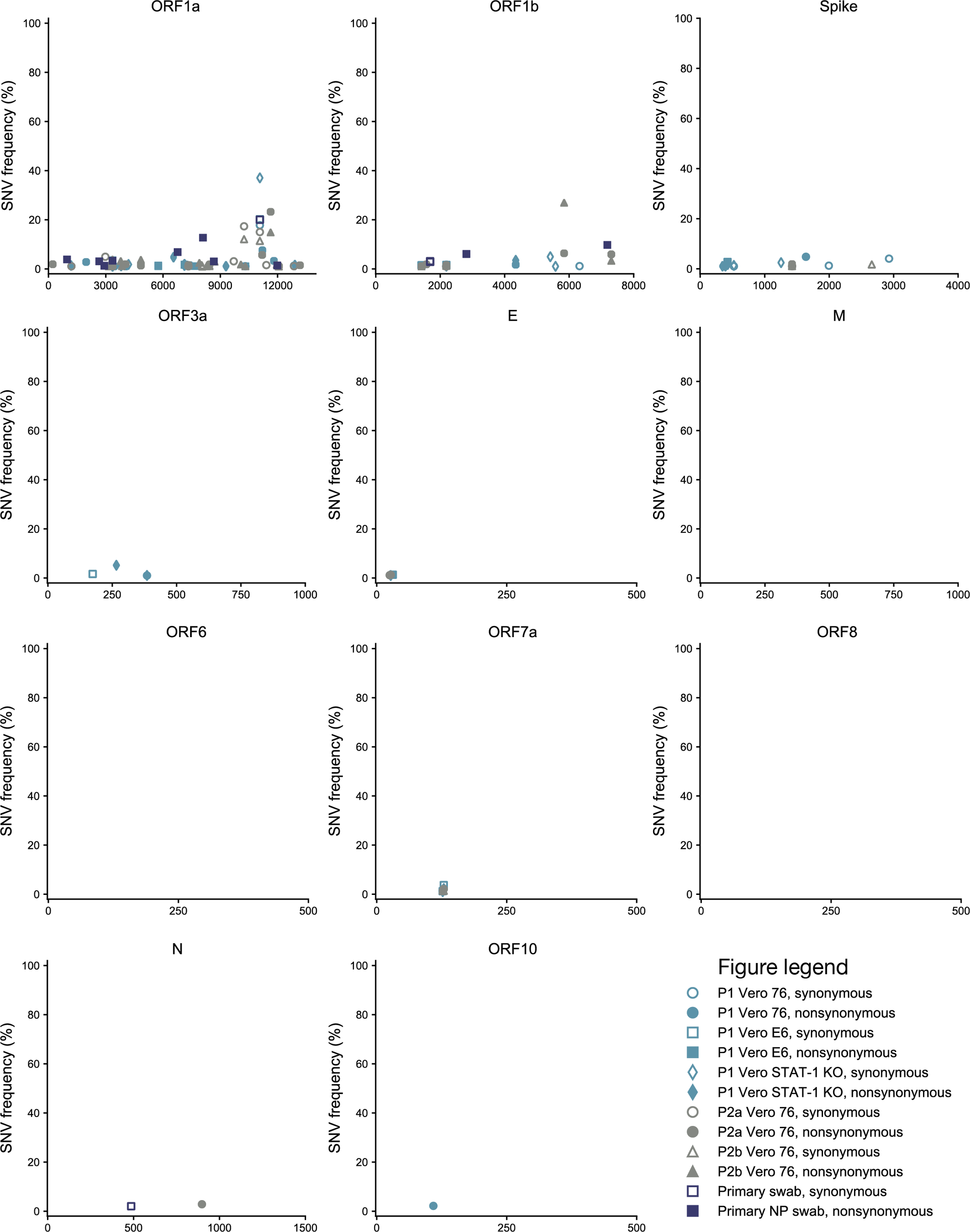
